# CD56 regulates human NK cell cytotoxicity through Pyk2

**DOI:** 10.1101/2020.03.19.998427

**Authors:** Justin T. Gunesch, Amera L. Dixon, Tasneem Ebrahim, Melissa Berrien-Elliott, Swetha Tatineni, Tejas Kumar, Everardo Hegewisch Solloa, Todd A. Fehniger, Emily M. Mace

## Abstract

Natural killer (NK) cells are innate immune cells that control viral infection and tumorigenic cell growth through targeted cell lysis and cytokine secretion. Human NK cells are classically defined as CD56^+^CD3^−^ in peripheral blood. CD56 is neural cell adhesion molecule (NCAM1), and despite its ubiquitous expression on human NK cells, the role of CD56 in human NK cell cytotoxic function has not been fully explored. In non-immune cells, NCAM can induce signaling, mediate adhesion, and promote exocytosis, in part through interactions with focal adhesion kinase (FAK). Here we describe the generation and use of CD56-deficient human NK cell lines to define a novel requirement for CD56 in target cell lysis. Namely, we demonstrate that deletion of CD56 on the NK92 cell line led to impaired cytotoxic function against multiple susceptible target cell lines. Deletion of CD56 in a second NK cell line, YTS cells, led to a less severe cytotoxicity defect but impairment in cytokine secretion. Confocal microscopy of wild-type and CD56-KO NK92 cells conjugated to susceptible targets revealed that CD56-KO cells failed to polarize during immunological synapse (IS) formation and had severely impaired exocytosis of lytic granules at the IS. Phosphorylation of the FAK family member Pyk2 at tyrosine 402 was decreased in NK92 CD56-KO cells, demonstrating a functional link between CD56 and IS formation and signaling in human NK cells. Cytotoxicity, lytic granule exocytosis, and the phosphorylation of Pyk2 were rescued by the reintroduction of NCAM140 (CD56), into NK92 CD56-KO cells. These data highlight a novel functional role for CD56 in stimulating exocytosis and promoting cytotoxicity in human NK cells.

## Introduction

Natural killer (NK) cells are innate immune effectors that play an important role in the clearance of virally infected and tumorigenic cells and modulation of immune responses. Human NK cells represent approximately 10% of circulating lymphocytes and can be defined within this population as CD56^+^CD3^−^ cells. CD56^dim^ cells are the predominant subset in peripheral blood while the minority of the circulating NK cell population are CD56^bright^ cells. CD56^bright^ and CD56^dim^ NK cells have unique expression of cell surface receptors, transcription factors and intracellular effector molecules that contribute to their distinct phenotypic and functional capacities (1, 2). Despite its conserved expression on human NK cells and its use as a phenotypic identifier, the role of CD56 in immune function is poorly defined.

CD56 is the cluster of differentiation nomenclature for neural cell adhesion molecule (NCAM). In addition to being abundantly expressed in cells of neuronal origin, NCAM is also expressed in other tissues including the heart, kidney, skeletal muscles, liver, and on hematopoietic-derived cells including dendritic cells, natural killer T (NKT) cells, and NK cells (3–5). NCAM is part of the immunoglobulin superfamily of cell adhesion molecules and has multiple isoforms due to RNA splicing (6–8). The three most abundant isoforms are the glycophosphatidylinositol (GPI)-anchored 120 kDa form (NCAM120) and two isoforms that contain transmembrane endodomains, NCAM140 and NCAM180 (6, 9). The extracellular portion of NCAM is highly conserved among the isoforms and contains 5 Ig-domains and two fibronectin-III domains that mediate homophilic (NCAM-NCAM) and heterophilic interactions between cells and the extracellular matrix (10, 11). NCAM^−/−^ mice have impaired learning and memory and behavioral disorders that are attributed to impaired nervous system development and post-differentiation maintenance of signaling and plasticity (12, 13). As such, NCAM plays an important role in non-immune cellular differentiation and function.

In addition to its role as an adhesion molecule, NCAM signaling in neuronal cells is important for neurite outgrowth and synaptic plasticity. NCAM140 is constitutively associated with the membrane-associated Src-family tyrosine kinase Fyn in axonal growth cones (14). Signaling through NCAM140 by agonist antibodies induces the recruitment of the non-receptor tyrosine kinase focal adhesion kinase (FAK) in neuronal cells, leading to a rise in intracellular Ca++ and neurite outgrowth (14–16). In addition to this signaling pathway, NCAM activates fibroblast growth factor receptor -1 (FGFR1) following trans-homophilic binding or binding of soluble extracellular NCAM and activates a non-canonical signaling pathway via phospholipase Cγ (PLCγ) that activates Erk1/2 in a Src-kinase dependent manner (17).

NCAM is unique amongst other glycoproteins in that its fifth Ig-domain contains 2 N-linked glycosylation sites that can be highly polysialylated (6, 9). PSA-NCAM is particularly important for synaptic plasticity and is highly expressed in embryonic development, with decreasing and restricted expression found in adults. The phenotype of olfactory bulb precursor cell migration in NCAM-deficient mice can be recapitulated by enzymatic removal of PSA, demonstrating the importance of PSA in NCAM signaling (18). The role of PSA addition to NCAM in NK cells has not been explored, and the lack of expression of NCAM or a known NCAM homologue on murine NK cells precludes the use of mouse models to test the in vivo requirement for NCAM or PSA-NCAM on NK cells.

NK cell cytotoxic function is exerted through the directed secretion of perforin- and granzyme-containing lytic granules following formation of an immunological synapse (IS) that includes adhesion, effector and termination stages (Orange, 2008). Cell adhesion is mediated by integrins, particularly LFA-1, that play a critical role in IS formation and actin polymerization. Prior to the polarization of the microtubule organizing center (MTOC) to the IS, lytic granules converge to the MTOC in a dynein-dependent minus-ended directed manner that is independent of actin polymerization and microtubule dynamics; as such, ligation of LFA-1 or CD28 alone is sufficient to induce convergence (19, 20). LFA-1-mediated outside-in signaling leads to F-actin polymerization and reorganization, phosphatidylinositol 4,5-bisphosphate (PtdIns(4,5)P2) generation, and the activation of protein tyrosine kinases, including Src family kinases (21, 22). Activating signaling and sustained F-actin remodeling promotes the polarization of the MTOC, with converged lytic granules, to the lytic IS (23–25). Ultimately, lytic granules transverse a dynamic yet pervasive cortical actin network to access the plasma membrane, where they undergo SNARE-mediated exocytosis at the IS (24, 26–28).

Previous studies have demonstrated that CD56 can promote human NK cell cytotoxic function against some CD56^+^ target cells (29–32), although in some cases lysis is not enhanced by target cell CD56 expression (4). As these studies focused on homotypic CD56-mediated interactions, the significance of CD56 binding to heterotypic ligands is unclear yet is likely relevant. In neuronal cells, CD56 binds FGFR1 in cis and in trans, and CD56 on NK cells binding to FGFR1 on T cells leads to IL-2 production (33). CD56 also mediates direct recognition of A. fumigatus leading to fungal-induced NK cell production of MIP-1α, MIP-1β and RANTES (34). This interaction is marked by accumulation of CD56 at the interface between the NK cell and A. fumigatus and is actin-dependent (34).

While CD56 has been implicated in NK cell development, migration and cytotoxicity (4, 23, 30, 31, 35), the signaling pathways that regulate its function in immune cells have not been described. Given signaling downstream of CD56 that is mediated by FAK in neuronal cells, one potential link between CD56 and IS formation is the closely related non-receptor tyrosine kinase 2 (Pyk2), which is highly expressed in NK cells (36). Pyk2 colocalizes with the MTOC in the uropod of migrating NK cells, however following activation it is translocated to the IS and is required for MTOC polarization in IL-2 activated primary NK cells (37). Expression of dominant negative Pyk2 disrupts cytotoxicity in this system, and its interactions with β1 integrin, paxillin and other protein tyrosine kinases suggests that Pyk2 plays a role as a scaffolding protein that helps orchestrate NK cell cytotoxicity (36–38). Here, we describe a requirement for CD56 in human NK cell function and show that deletion of CD56 in two human NK cell lines leads to impaired secretion and accompanying lytic dysfunction. Furthermore, we identify Pyk2 as a critical signaling intermediate downstream of CD56. These data demonstrate a direct role for CD56 in the NK cell-mediated lysis of CD56-negative target cells and describe a novel activation pathway for cytotoxicity that is unique to human NK cells.

## Results

### Characterization of CD56 expression and polysialation in primary cells and NK cell lines

We previously used CRISPR-Cas9 to generate stable CD56-knockout (KO) NK92 cell lines and define a requirement for CD56 in human NK cell migration (35). To extend our findings to a second NK cell line, we generated YTS CD56-KO cell lines using the same approach and CRISPR guides. CD56-negative YTS cells were isolated by FACS and the absence of CD56 protein was confirmed in both YTS and NK92 CD56-KO cell lines by Western blot analysis and flow cytometry (Fig 1A, B).

**Fig. 1.**
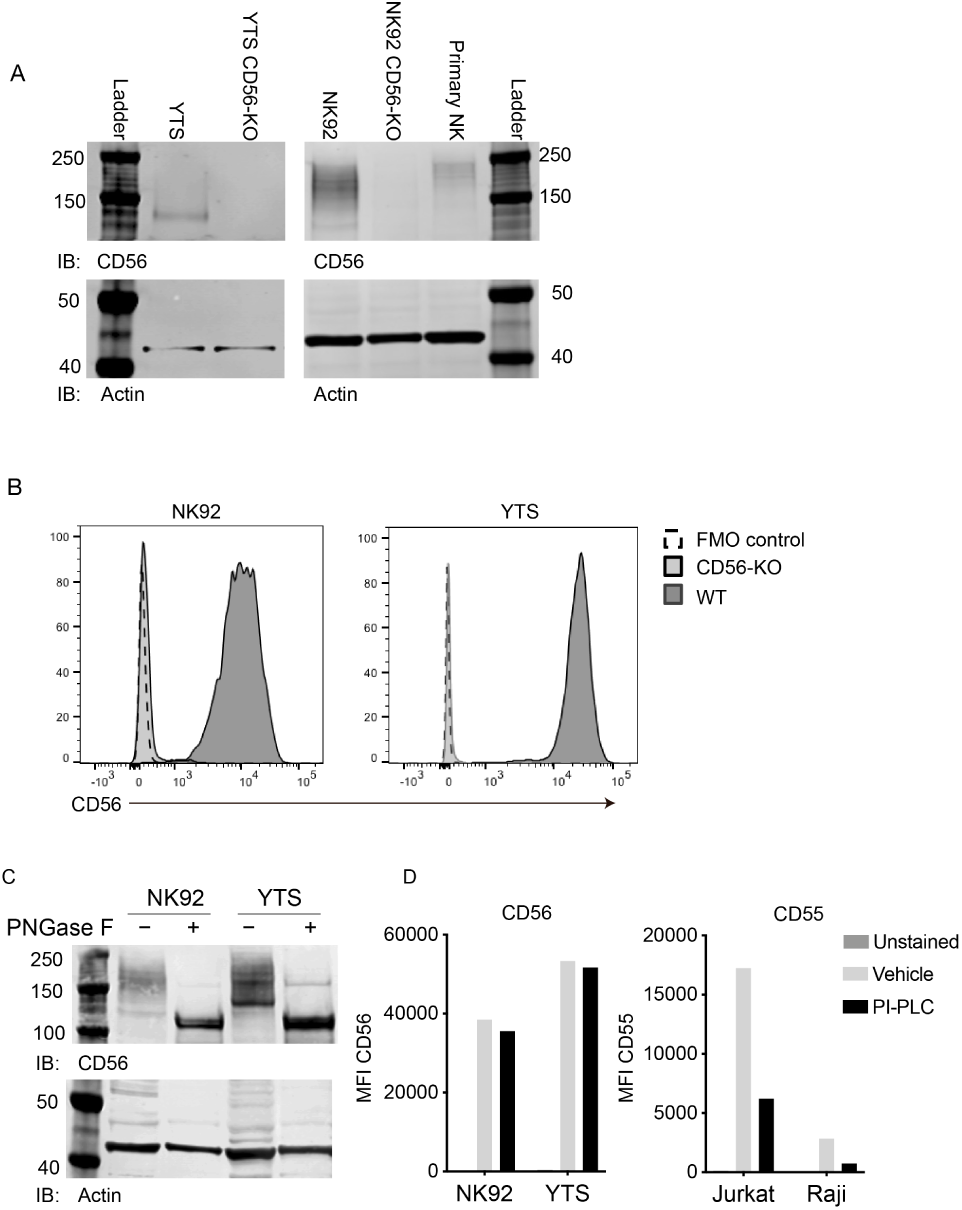
Validation of CD56 deletion in human NK cell lines and characterization of CD56 and its polysialation in human NK cells. A) Western blot analysis of CD56 from wild-type (WT) and CD56-knockout (KO) NK92 (left) and YTS (right) cell lines or primary human NK cells with actin as a loading control. B) Flow cytometry analysis of CD56 expression in NK92 or YTS WT (filled histogram, dark grey) or CD56-KO (filled histogram, light grey) cells compared to unstained cells (dashed line). C) NK92 or YTS cells were treated with PNGase F to remove polysialic acid. Following treatment, lysates were separated by SDS-PAGE and CD56 or actin as a loading control was detected by Western blotting. D) NK cell lines (left) or Jurkat or Raji cells as a positive control (right) were treated with PI-PLC to cleave GPI anchored proteins from the cell surface. PI-PLC activity was confirmed by cleavage of GPI-anchored CD55 (right) .xs

While transcripts for all three commonly expressed NCAM isoforms (NCAM140, NCAM180, NCAM120) can be detected in human NK cells, NCAM140 has been previously reported to be the only isoform expressed (4, 39). The extracellular domain of NCAM can be also post-translationally modified by the addition of polysialic acid (PSA), which affects the molecular weight of CD56 when detected by Western blotting. We noted that Western blot analyses of NK92 and YTS cell lines suggested that CD56 is highly polysialated, particularly in the NK92 cell line, leading to a range of apparent molecular weights including a band corresponding to the 140 kDa isoform (Fig. 1A). In contrast, the YTS cell line expressed primarily the 140 kDa isoform with less polysialation, whereas primary NK cells expressed higher molecular weight isoforms and, as previously reported, also expressed polysialated CD56 (40). To determine the contribution of polysialation to the molecular weight of CD56 on YTS and NK92 cell lines, we performed enzymatic treatment of cell lysates with PNGase F to cleave polysialic acid followed by Western blotting with anti-CD56 antibody. These data showed that removal of PSA reduced the variability of CD56 molecular weights and suggested that the 140 kDa or 120 kDa isoform were predominantly expressed in NK cell lines (Fig. 1C). Similar treatment followed by immunoblotting with a PSA-NCAM-specific antibody demonstrated that PSA-NCAM was not detectable following PNGase F treatment (Supp. Fig. 1), thus confirming that the treatment was highly effective in PSA removal.

Despite the reduction in molecular weight variability generated by enzyme treatment, it was difficult to define whether the most predominant band in the cell lines was the 120 kDa or 140 kDa isoform. To determine whether NCAM120, which is GPI-anchored, was expressed on the cell surface of NK cell lines, we treated cells with phosphoinositide phospholipase C (PI-PLC) and quantified expression of NCAM on the cell surface by flow cytometry. CD56 expression on WT NK92 and YTS cells was resistant to PI-PLC cleavage while CD55, a GPI-anchored protein, present on Raji and Jurkat cells, was cleaved (Fig. 1D). Together, these data strongly suggest that the isoform expressed on the surface of NK92 and YTS cells is not GPI-anchored NCAM120, but is primarily NCAM140 as previously reported (4, 9).

### CD56-KO NK cells have impaired lytic function towards CD56-negative targets

Previous reports on the role of CD56 in NK cell lytic function have focused on homotypic interactions between CD56 on NK cells and target cells and have led to conflicting conclusions about the role that CD56 plays in cytotoxicity (4, 30–32). To test the cytotoxic function of wild-type (WT) and CD56-knockout (KO) cell lines against CD56-negative target cells, we performed ^51^Cr-release cytotoxicity assays. NK92 or YTS cells were used as effectors against K562 target cells, which are susceptible to NK92-mediated lysis, or 721.221, which are susceptible to lysis by both cell lines (41). Deletion of CD56 in the NK92 cell line severely abrogated its cytolytic function against susceptible K562 and 721.221 target cell lines (Fig. 2A). However, YTS CD56-KO cells exerted normal lytic function against 721.221 target cells when compared to their WT counterparts (Fig. 2B). To test a second susceptible target with YTS cells, we used KT86 cells, which are K562 cells that express CD86 and thus are susceptible to YTS-mediated lysis (42). The use of KT86 targets did not reveal a requirement for CD56 in their lysis by YTS cells despite the failure of NK92 CD56-KO cells to lyse K562 targets (Fig. 2B). These data demonstrated the conserved defect in NK92 CD56-KO cytolytic function, whereas YTS cells were not significantly affected by loss of CD56 in their killing of multiple target cell lines in a 4-hour ^51^Cr assay.

**Fig. 2.**
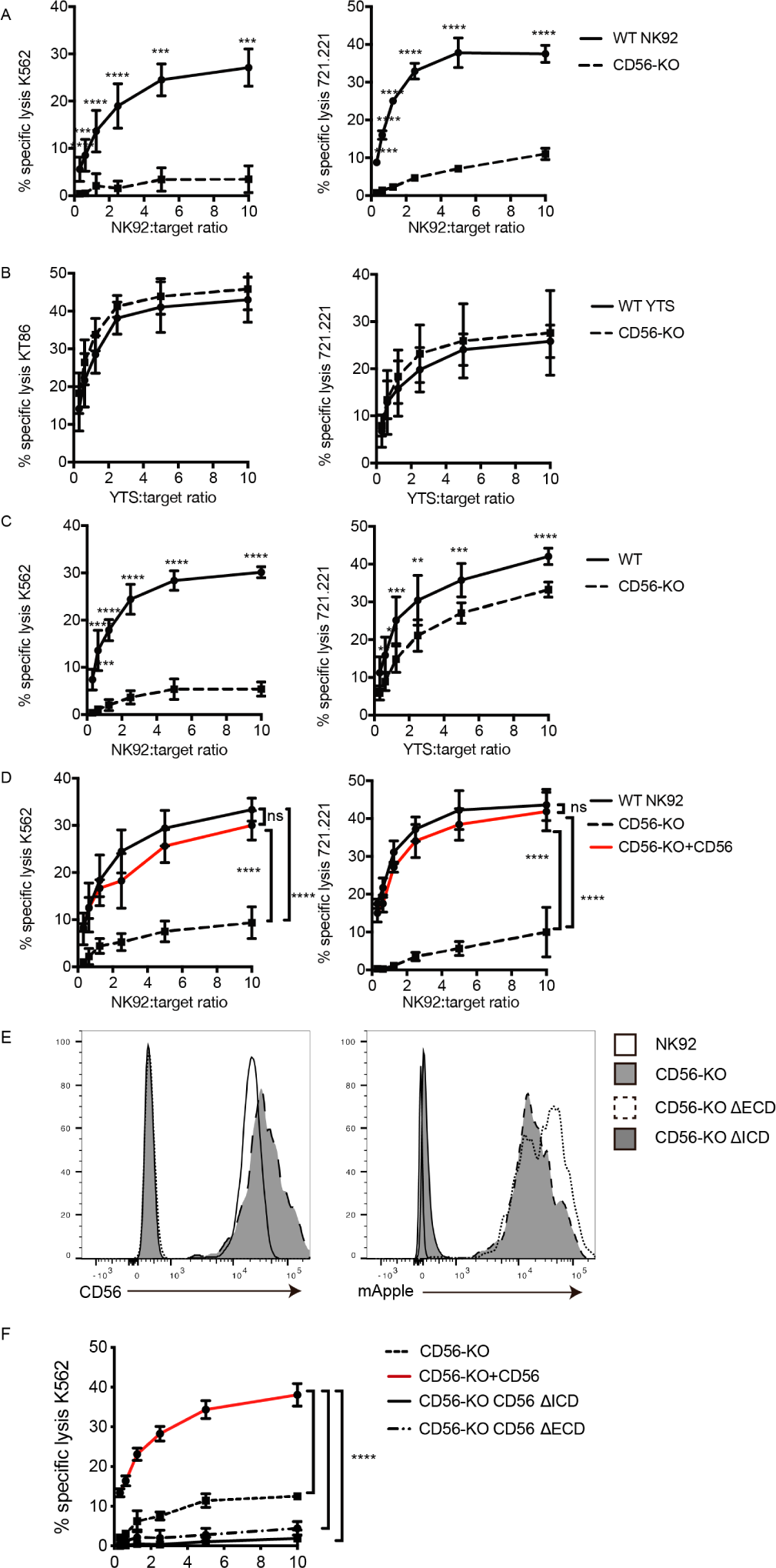
CD56 deletion abrogates NK92 cytotoxic function and delays YTS cytotoxicity. ^51^ Cr-release assays were performed using NK92 (A, C, D) or YTS (B) WT and CD56-KO cell lines as effectors against susceptible targets. A) 4-hour assays were performed with NK92 cell lines using K562 (left) or 721.221 (right) target cells. B) 4-hour assays were performed with YTS cell lines using KT86 (left) or 721.221 (right) target cells. C) 1-hour ^51^ Cr-release assays were performed using NK92 (left) or YTS (right) cells as effectors. D) CD56 (NCAM-140) was re-expressed in NK92 CD56-KO cells and these cells or NK92 CD56-KO cells were used for 4-hour cytotoxicity assays against K562 (left) or 721.221 (right) target cells. E) CD56-KO NK92 cells were transfected with chimeric CD56 constructs fused to an mApple fluorescent reporter as described in Methods. Flow cytometry was used to confirm the expression of CD56 and/or mApple. F) Cytotoxicity assays were performed with chimeric cell lines using K562 cells as targets. Mean±S.D of at least three independent experiments pooled. **p<0.01, ***p<0.001, ****p<0.0001 by Ordinary one-way ANOVA with multiple corrections test or unpaired student t-test with Welch’s correction. ΔECD: chimeric construct lacking extracellular domain, ΔECD: chimeric construct lacking intracellular domain.

NK cell cytotoxic function includes serial killing of target cells, with target cell lysis followed by NK cell detachment and re-engagement with subsequent targets (43–46). Given the demonstrated defect in NK cell migration in CD56-KO NK92 cells (35), we sought to measure whether the defect in NK cell killing could be attributed to impaired serial killing. The average time to first kill of a target cell by an NK92 or YTS cell ranges from 35-45 minutes (41), therefore to test the efficacy of early cytotoxic function we performed ^51^Cr cytotoxicity assays with 1 hour incubations with target cells. As seen in 4-hour assays, the lytic function of NK92 CD56-KO cells was severely impaired following 1 hour of incubation with target cells, suggesting that the decrease seen at 4 hours is not due to defects in serial killing. In addition, 1-hour ^51^Cr-release assays with YTS CD56-KO cells revealed a decreased specific lysis compared to WT YTS (Fig. 2C). While the killing decrease in the YTS CD56-KO cells at 1 hour was not as substantive as observed in NK92 CD56-KO cells, it was statistically significant at all effector to target ratios and consistent between multiple experiments. Therefore, while the impairment in CD56-deficient NK92 cells was more profound, there was a conserved phenotype between the two human NK cell lines tested.

To determine whether the observed defect in cytotoxicity was due to dysregulated expression of molecules associated with NK cell lytic function, multiparametric flow cytometry was performed using 5 panels designed to measure expression of receptors required for NK cell development, adhesion, activation, and inhibition (47). Small increases in the percent positive cells and mean fluorescence intensity of CD2, CD11a, CD18, CD45, CD94, and NKG2A were observed in NK92 CD56-KO cells (Supp. Fig. 2). However, these slight differences may be attributed to better ligand accessibility by antibodies, as deleting CD56 removes the long chains of negatively charged polysialic acid attached to CD56 (48). No significant differences were observed in the frequency of NK92 CD56-KO cells that were positive for expression of granzyme A or B and for IFNγ at rest or after activation with PMA/ionomycin relative to the parental NK92 cell line (Supp. Fig. 2). Similarly, we did not observe differences in surface receptors or effector molecules at rest or after activation in YTS CD56-KO cells when compared to wild-type YTS (Supp. Fig. 2). Therefore, we concluded that the defects in cytotoxicity that we observed were not due to altered expression of necessary activating receptors or effector molecules on the NK92 cell line.

To confirm that the defect in NK92 cytotoxicity was specifically conferred by CD56 deletion, NK92 CD56-KO cells were retrovirally transduced with the NCAM140 isoform fused with an mApple fluorescent reporter to re-introduce CD56 expression. CD56 protein expression and expression of polysialic acid were confirmed in the reconstituted cells by flow cytometry (Supp. Fig. 3). Importantly, re-expression of CD56 restored the lytic function of NK92 CD56-KO cells against both 721.221 and K562 targets (Fig. 2D), demonstrating that the impairment we observed was specific to the deletion of CD56 and that re-expression of NCAM140 was sufficient to rescue this defect.

We sought to further define the requirements for the intra-cellular and extracellular domains of CD56 in NK cell cytotoxicity. The 140 kDa isoform of CD56 (NCAM) expresses 5 Ig-like domains and 2 FNIII repeats, a transmembrane domain and an intracellular domain. We generated CD56 chimeric constructs and retrovirally transduced these into NK92 CD56-KO cells. ΔICD lacks the intracellular domain (ICD) of CD56 but contains the extracellular domain (ECD) and transmembrane domain (TM); ΔECD lacks the CD56 ECD but includes the transmembrane and intracellular domains. Stable NK92 CD56-KO cell lines were generated and confirmed to be mApple positive and the ΔICD cells were verified to be CD56^+^ (Fig 2E). ^51^Cr-release assays were performed to determine the requirements for CD56 domains in cytotoxicity in NK92 cells. Sole expression of either the ECD or ICD was not sufficient to rescue cytotoxicity (Fig. 2F), whereas full-length CD56 restored cytotoxicity as previously demonstrated (Fig. 2D, F). Collectively, these data indicate that the full-length NCAM140 protein is required to promote human NK cell cytotoxicity and suggests a role for intracellular interactions that mediate this process.

### Secretory function is decreased in NK92 and YTS CD56-KO cell lines

Both NK92 and YTS cell lines kill target cells via perforin- and granzyme-dependent exocytosis. In addition, YTS in particular are potent producers of IFNγ upon stimulation (49). To further assess the effect of CD56 deletion on NK cell function, two methods of detection were used; a colorimetric benzyloxycarbonyl-L-lysine thiobenzyl ester (BLT) esterase assay for detection of granzyme A function as a readout for granule exocytosis and detection of surface-exposed lysosomal-associated membrane protein-1 (LAMP-1 or CD107a) by flow cytometry after plate-bound activation or coculture with target cells (50–52). We activated WT, CD56-KO, and CD56 rescued CD56-KO NK92 cells for 90 minutes with immobilized anti-CD18 and NKp30 antibodies and measured BLT esterase activity (28). As suggested by the impaired lytic function demonstrated by ^51^Cr assays, NK92 CD56-KO cells had significantly decreased granzyme A activity compared to WT NK92, demonstrating that degranulation is impaired in the CD56 knockout cells. Re-expression of full-length CD56 in NK92 CD56-KO cells restored granzyme A activity comparable to that of WT NK92 (Fig. 3A).

**Fig. 3.**
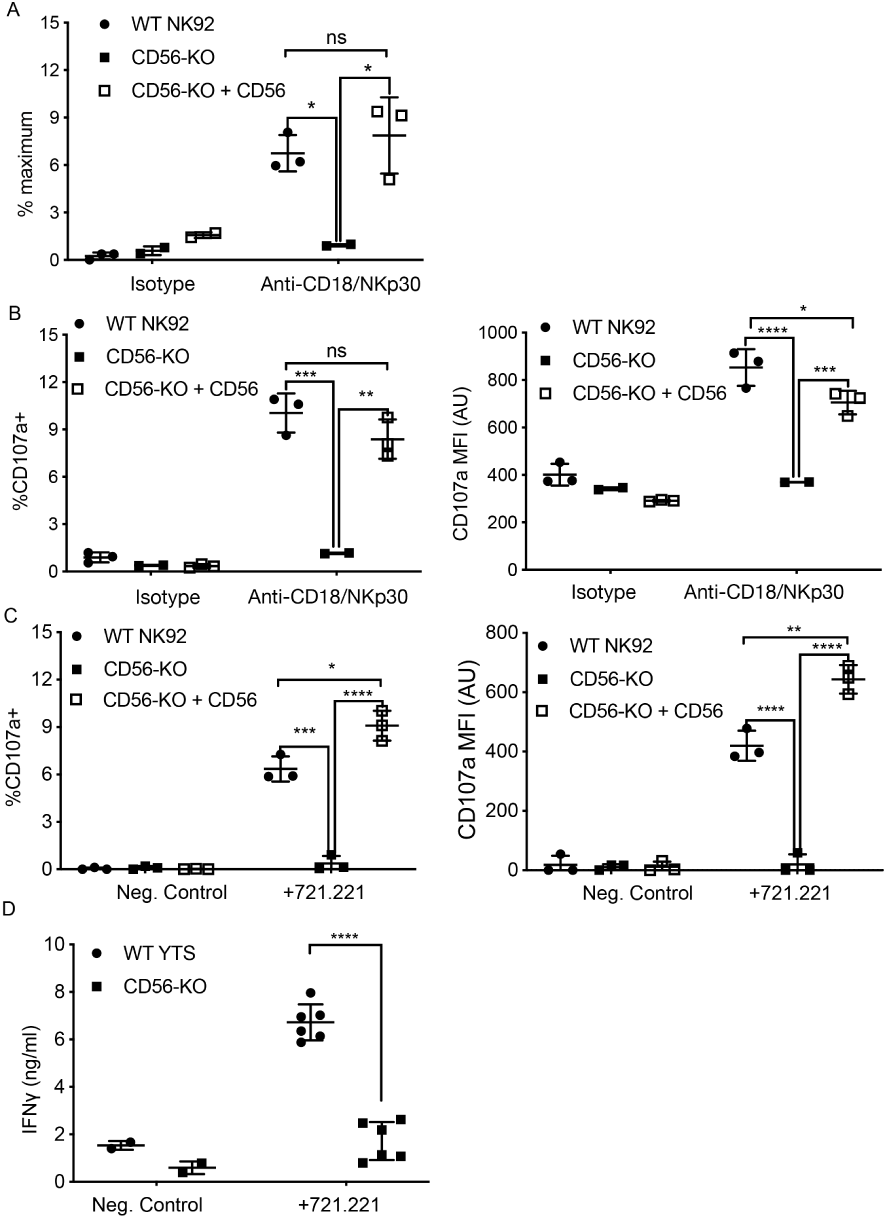
CD56 expression is required for exocytosis of NK92 cells. A) WT NK92, CD56-KO or CD56-KO reconstituted cells were incubated for 60-90 minutes on plates pre-coated with 10 µg/ml of anti-CD18 and anti-NKp30 antibodies. Supernatant was collected and granzyme A secretion was measured by a BLT esterase assay. Secretory potential was measured as a readout of the % maximum of granzyme A activity in the supernatant. B) WT NK92, CD56-KO or CD56-KO reconstituted cells were incubated for 1-2 hours on plates pre-coated with 10 µg/ml of anti-CD18 and anti-NKp30 antibodies. Cells were harvested and degranulation was measured by CD107a expression using flow cytometry. C) WT NK92, CD56-KO or CD56-KO reconstituted cells were co-cultured with 721.221 target cells. Cells were harvested and CD107a expression was measured by flow cytometry. For co-culture experiments the average background of media only was subtracted from samples. Mean ± SD. *p<0.05, **p<0.01, ***p<0.001, ****p<0.0001 by one-way ANOVA with Tukey’s multiple comparisons post-hoc test. D) WT YTS and CD56-KO cells were incubated with 721.221 target cells at a 2:1 ratio for 22 hours. Supernatant was collected and used in a human IFN gamma ELISA. ****p<0.0001 by unpaired student t-test with Welch’s correction. All data are representative of 3 independent experiments performed in duplicate or triplicate.

To further support our observation that CD56-KO NK92 cells did not undergo lytic granule exocytosis, we measured CD107a exposure following activation by plate-bound antibodies or target cells. Following plate-bound stimulation, NK92 CD56-KO had a significantly reduced percentage of cells positive for CD107a compared to WT NK92, and those cells that did express CD107a did so with a lower mean fluorescent intensity, supporting our data showing decreased secretion in the absence of CD56 (Fig. 3B). As with the BLT esterase assay, re-expression of CD56 in the knockout cells restored their capacity to degranulate (Fig. 3B). Furthermore, the same effect was observed when NK92 cells were co-incubated with target cells (Fig. 3C). Taken together, these data demonstrated that the deletion of CD56 in NK92 cells leads to a defect in lytic granule exocytosis in response to target cell activation or activating receptor cross-linking.

We failed to detect BLT esterase secreted from YTS cell lines, likely due to their very low expression of granzyme A (49). Furthermore, YTS cells undergo fewer degranulation events on a per cell basis when killing target cells than NK92 (41). Measurement of exposed CD10^7^a on YTS cell lines (WT and CD56-KO) after plate-bound activation proved difficult and inconclusive after coculture with 721.221 target cells (data not shown). However, YTS cells robustly produce IFNγ in response to activation (49). Co-incubation of WT YTS with 721.221 target cells led to secretion of IFNγ detectable by ELISA, whereas this secretion was significantly decreased in the CD56-KO YTS cells (Fig. 3D). We confirmed robust IFNγ production in YTS CD56-KO cells in response to stimulation with PMA/ionomycin by intracellular flow cytometry (Supp. Fig. 2), confirming that this decrease in IFNy levels from YTS cells conjugated with Y21.221 is due to a defect in secretion, rather than production. Therefore, despite their intact target cell killing in 4-hour ^51^Cr assays, YTS cells had significant impairment in secretion in response to contact-dependent activation when cytokine production was measured.

In summary, deletion of CD56 in the NK92 and YTS cell lines impairs their secretion in response to activation, including degranulation in NK92 CD56-KO cells, demonstrating a critical role for CD56 in NK function that is independent from CD56 homotypic interactions. The differential functional response to CD56 deletion of these cell lines supports their differing properties and speaks to relevant differences in their biology (49); here we have chosen to focus on the mechanism by which CD56 is mediating lytic function in the NK92 cell line.

### Lytic immune synapse formation and function

NK cell lytic function is exerted through the formation of an immunological synapse, which serves to focus directed secretion and mitigate bystander killing. The steps leading to IS formation can broadly be defined by adhesion/activation, polarization, secretion and termination stages (53). Initial activation through ITAM-based activating receptors, integrins, or cytokine stimulation leads to lytic granule convergence to the MTOC, a step that precedes actin remodeling and the re-orientation of the MTOC to the synapse and ultimately lytic granule exocytosis and target cell death.

Having shown that the secretion stage was impacted by CD56 deletion, we sought to further define how activation and polarization were affected in NK92 CD56-KO NK cell lines. Fixed cell confocal microscopy was performed to visualize granule convergence, MTOC polarization, and actin accumulation at the cell-cell interface and each of these parameters was quantified as previously described (54, 55). Both WT and CD56-KO NK92 cells were found in conjugates with K562 target cells, suggesting that their ability to adhere to targets was not impaired (Fig. 4A). While actin remodeling was consistently decreased in CD56-KO cells relative to WT NK92, this effect was not significant (Fig. 4B).

**Fig. 4.**
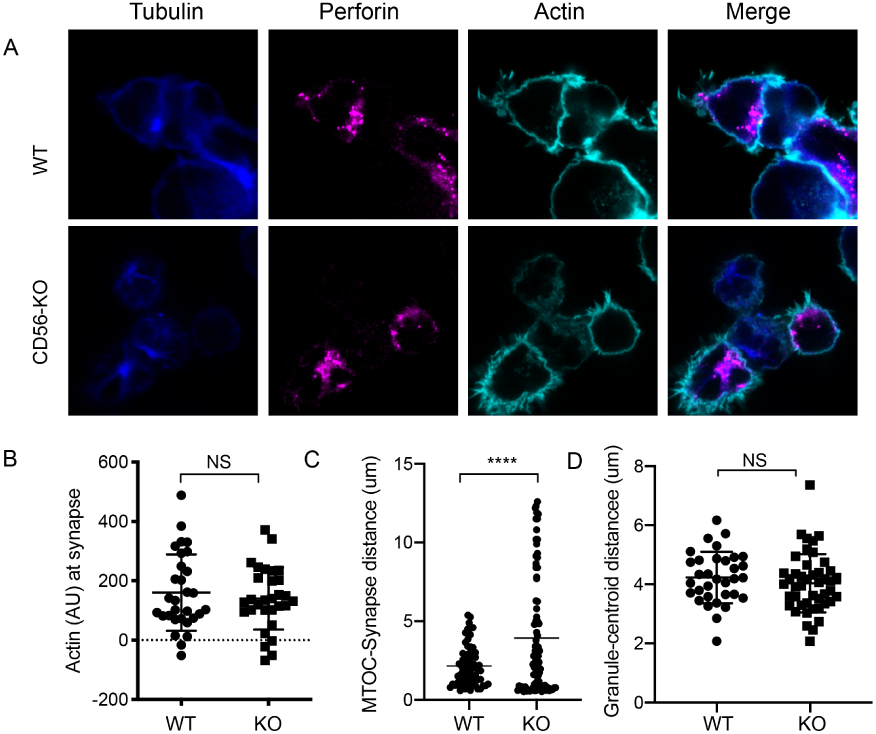
Impaired immune synapse formation in NK92 CD56-KO cells. WT or CD56-KO effector cells were cultured at a 2:1 ratio with K562 target cells for 45 minutes then fixed, immunostained and visualized by confocal microscopy. A) Representative images from >30 cells from >3 independent experiments immunostained as indicated. B) Integrated intensity of actin at the immune synapse for WT or CD56-KO cells. Data are representative of one experiment performed >3 times. n=30, 39. NS = not significant by unpaired t-test. C) MTOC to synapse distance (µm) calculated from WT or CD56-KO conjugates. n = 70, 83. Data pooled from 3 independent repeated experiments. D) Mean granule to centroid distance (MGD) for WT or CD56-KO conjugates. Each data point represents MGD from one conjugate. n= 33, 45 from one representative experiment of >3 experiments. NS = not significant by unpaired T test with Welch’s correction.

In contrast, the MTOC was not polarized towards the immunological synapse in CD56-KO NK92 cells as it was in NK92 cells, suggesting decreased activation. This observation was quantified by significantly greater MTOC-synapse distance (Fig. 4C). Therefore, loss of CD56 expression in the NK92 cell line, which results in impaired cytotoxic function, is accompanied by impaired formation of the immunological synapse reflected by impaired MTOC polarization towards target cells.

### Pyk2 colocalizes with CD56 and its phosphorylation is decreased in NK92 CD56-KO cells

The introduction of the CD56-domain specific constructs into the NK92 CD56-KO cells illustrated the observation that both the extracellular and intracellular domain of CD56 are required to recover cytotoxicity. This suggests that the intracellular domain of CD56 may be interacting with intracellular molecules within NK92 cells to enable cytotoxic function. Protein tyrosine kinase 2 (Pyk2) is closely related to focal adhesion kinase (FAK) and a predicted interacting partner of CD56 through binding to Fyn (Sancho et al., 2000; Szklarczyk et al., 2015). In non-immune cells, engagement of NCAM140 (CD56) recruits and activates FAK, which in turn activates the MAPK signaling pathway that is important for neurite outgrowth and cell survival (Beggs et al., 1997; Ditlevsen et al., 2003; Schmid et al., 1999). Additionally, Pyk2 is an important regulator of NK cell cytotoxicity by stimulating lytic granule polarization and target cell conjugation (Gismondi et al., 2000; Sancho et al., 2000; Zhang et al., 2014). To determine the effect of CD56 deletion on Pyk2 phosphorylation WT, CD56-KO, and CD56-KO reconstituted NK92 cells were activated by plate-bound anti-NKp30 and -CD18. Cells were dissociated and phosphorylation of Pyk2 (Y402) was measured by intracellular flow cytometry. We found that phosphorylation of Pyk2 tY402 was significantly decreased in the NK92 CD56-KO cells, a phenotype that was recovered by the reconstitution of CD56 (Fig. 5A).

**Fig. 5.**
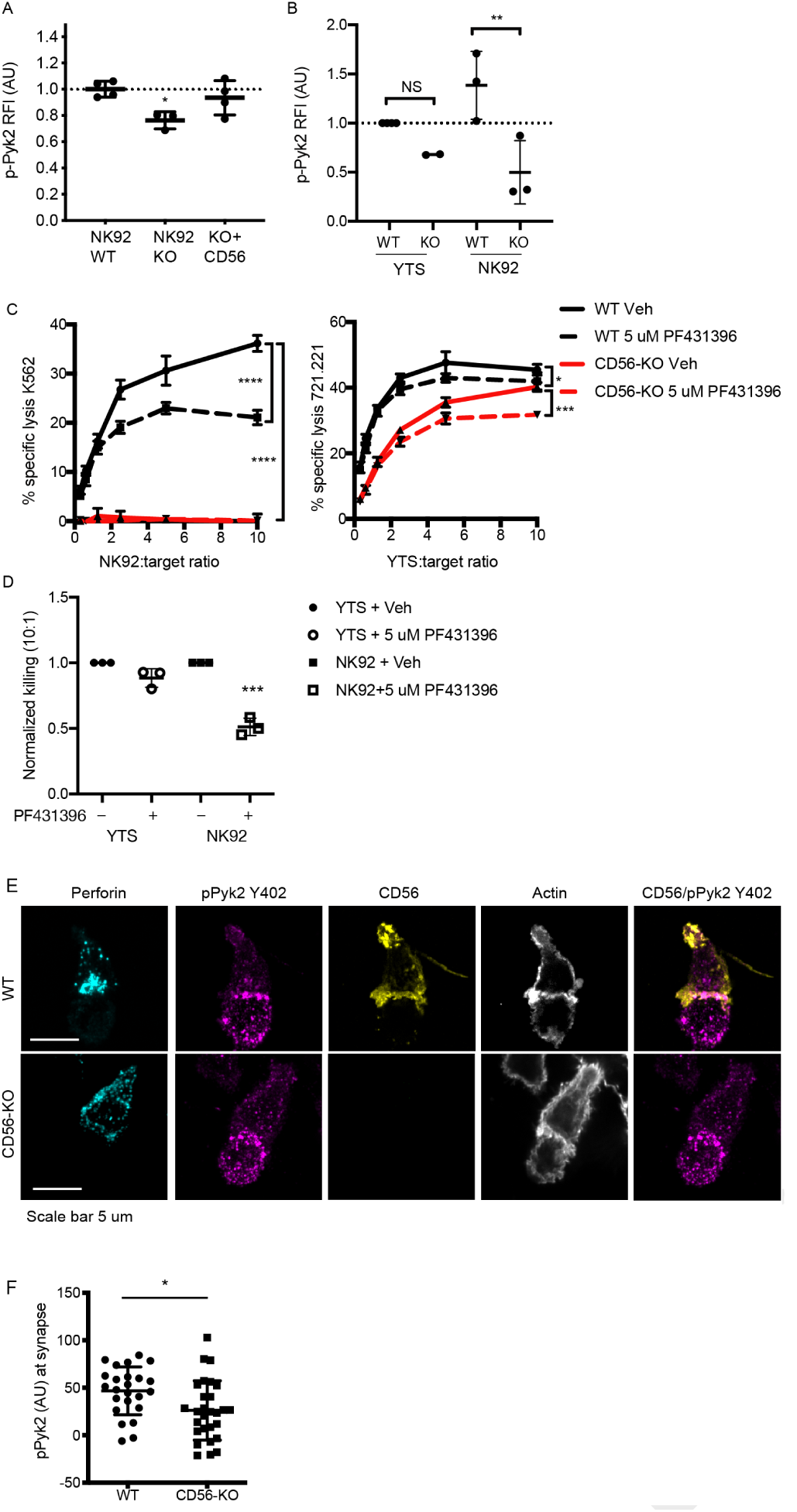
Phosphorylation of Pyk2 is decreased in NK92 CD56-KO cells upon activating receptor ligation. A) NK92 WT, CD56-KO or CD56 reconstituted (KO+CD56) cells were incubated for 25-30 minutes on plates pre-coated with 10 µg/ml of anti-CD18 and anti-NKp30 antibodies. Cells were permeabilized and immunostained for p-Pyk2 Y402 then data were acquired by flow cytometry. Relative fluorescent intensity (RFI) of pPyk2 was calculated based upon the intensity of the WT NK92 condition. Shown are the pooled data from 3 independent experiments. B) WT or CD56-KO NK92 or YTS cells were permeabilized and immunostained for pPyk2 Y402 then data were acquired by flow cytometry. Shown is pooled data from 2 (YTS) or 3 (NK92) independent experiments. **p<0.01 by one-way ANOVA with multiple comparisons. C) 4 hour ^51^ Cr assays were performed with WT (black) or CD56-KO (red) NK92 cells as effectors. Assays were performed in the presence of Pyk2 inhibitor PF431396 or vehicle control (DMSO) following brief pre-incubation of effectors with inhibitor. Shown are representative data from 3 independent repeats. D) Pooled data from the 10:1 effector to target cell ratio of the experiments described in (C) normalized to the WT YTS condition without inhibitor. ***p<0.001 by one-way ANOVA with multiple comparisons. E) Representative confocal microscopy images of WT or CD56-KO NK92 effectors conjugated to K562 target cells in the presence of non-blocking CD56 antibody then fixed and immunostained for perforin, pPyk2 Y402 and actin. F) Fluorescent intensity of pPYK2 Y402 at the immune synapse of WT or CD56-KO effector cells. n = 24, 28 from one representative experiment of >3 independent repeats.

We hypothesized that the differential impact upon NK cell function on YTS and NK92 cell lines following CD56 deletion may be related to differential usage of Pyk2-dependent signaling pathways. To test this hypothesis, we analyzed Pyk2 phosphorylation in YTS and NK92 WT and CD56-KO cells using flow cytometry as described above. While we noted a similar trend in YTS cells, notably decreased Pyk2 phosphorylation in CD56-KO cells relative to WT, this effect was not as profound as that seen in NK92 cells and was not statistically significant (Fig. 5B). In addition, the magnitude of Pyk2 phosphorylation in WT YTS cells was lower than that of NK92 cells based upon normalized fluorescence intensities. The lower relative fluorescence intensity of Pyk2 Y402 phosphorylation in YTS cells and greater relative impairment in NK92 CD56-KO cells suggests a greater dependence on Pyk2 phosphorylation in NK92 cells and a potential mechanism whereby CD56 deletion has a greater effect on Pyk2-mediated cytotoxic function in NK92 than YTS cells. To further investigate this hypothesis, we performed ^51^Cr assays in the presence of PF431396, a Pyk2/FAK inhibitor that inhibits Pyk2 autophosphorylation and blocks its kinase function (56). Consistent with the differential phosphorylation of CD56 in YTS and NK92 cells, the presence of Pyk2 inhibitor significantly decreased the cytotoxic function of WT NK92 cells, whereas both WT and CD56-KO YTS cells were only minimally affected (Fig. 5C, D). These data demonstrate that, in the absence of CD56 expression in NK92 cells, Pyk2 phosphorylation is decreased and that inhibition of Pyk2 phosphorylation in NK92 cells impairs cellular cytotoxicity. In YTS cells, a reduced dependence on Pyk2 for target cell lysis is reflected by retained lytic function both in the presence of Pyk2 inhibitor and the absence of CD56-mediated function.

We further sought to determine whether Pyk2 localization at the immune synapse was affected by loss of CD56. Fixed cell confocal microscopy was performed by acquiring 3D volumes of WT or CD56-KO NK92 cells conjugated to K562 target cells and immunostained for pPyk2 Y402, perforin and actin. As suggested by intracellular flow cytometric analysis of Pyk2 phosphorylation, we detected higher fluorescence intensity of pPyk2 Y402 in WT NK92 relative to CD56-KO NK92 (Fig. 5E). Quantification of conjugates confirmed greater pPyk2 accumulation at the IS of WT NK92 cells when compared to CD56-KO NK92 cells (Fig. 5F). In contrast, we measured no difference in the localization or intensity of total Pyk2 between WT and CD56-KO cell lines, which primarily localized to the MTOC as previously reported (37) (Supp. Fig. 4). These measurements were made difficult by the significant phosphorylation of Pyk2 detected in the target cells, however the effect of CD56 deletion on Pyk2 phosphorylation is underscored by our functional experiments in a target cell-free system (Fig. 5A).

Both Pyk2 and CD56 are localized to the uropod of migrating human NK cells (Mace et al., 2016; Sancho et al., 2000). We noted that CD56 in NK92 cells conjugated to targets remained partially localized to the uropod, however a fraction of CD56 localized to the IS, where it strongly co-localized with actin and pPyk2 Y402 (Fig. 5E). This was observed both early and late in immune synapse formation (data not shown), suggesting that the effect we observed was not due to the kinetics of CD56 re-distribution. These data support our functional studies demonstrating that loss of CD56 in NK92 cells impairs immune synapse function through deregulated Pyk2 activation, while additionally illustrating the recruitment of CD56 to the IS in human NK cells.

### Cytotoxic function of CD56-deficient primary NK cells is intact

We sought to determine the effect of CD56 deletion on primary NK cell function. Using CRISPR-Cas9, we deleted CD56 in bulk peripheral blood NK cells or CD56^bright^ NK cells activated with IL-15. CD56 expression was reduced by 9-fold when cells were incubated with low dose IL-15 but was still detectable on the surface of NK cells (data not shown). Because CD56 protein appeared stable, we utilized higher doses of IL-15 to induce NK cell proliferation and CD56 dilution. Using this approach, the deletion of CD56 was highly efficient following delivery and expansion with IL-15 (Fig. 6A); notably, NK cell cytotoxic function was intact (Fig. 6B).

**Fig. 6.**
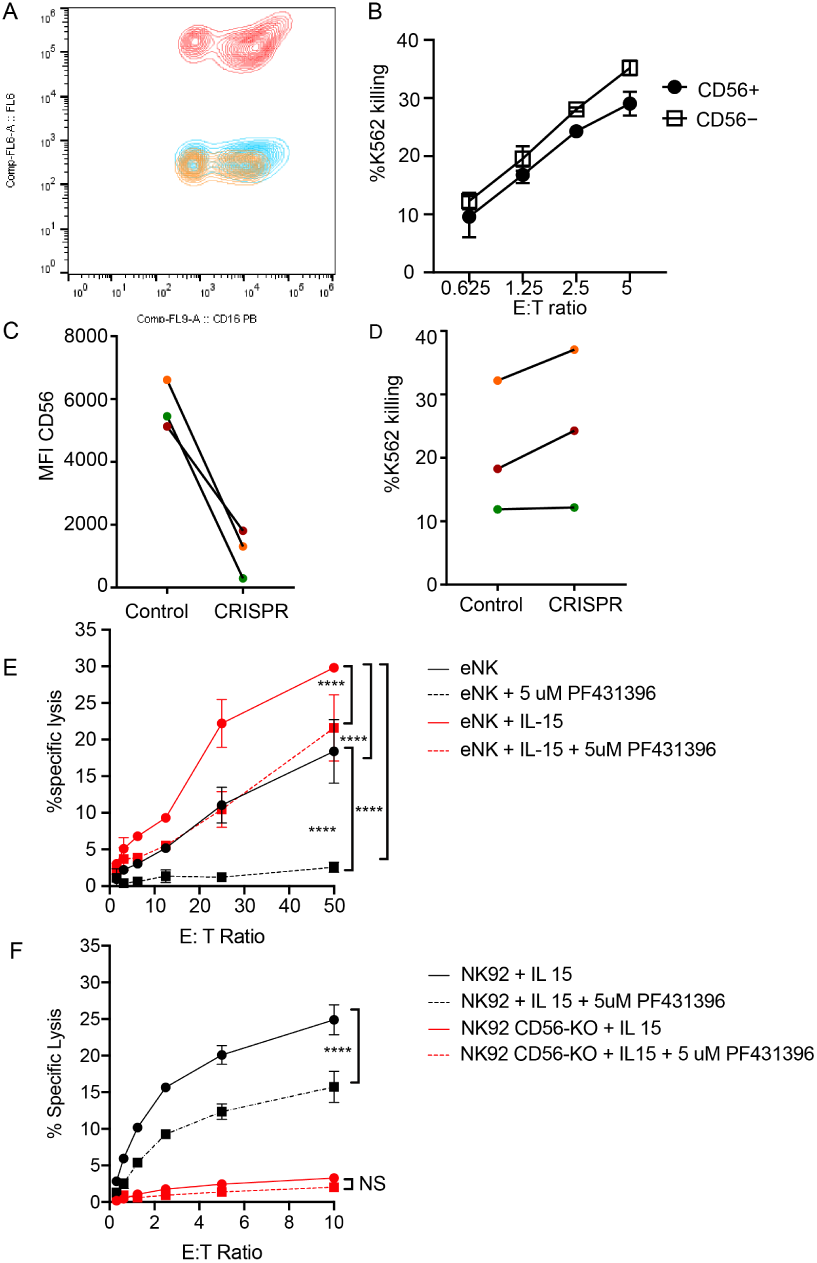
CD56-deficient primary NK cells retain lytic function. Primary NK cells were isolated and allowed to rest overnight in the presence of low-dose IL-15 prior to delivery of CD56 CRISPR-Cas9. Cells were further expanded in the presence of 25 ng/ml IL-15 for 15 days and cytotoxicity against K562 targets was measured. A) Representative FACS plot of CD56-deficient (blue) or control primary cells (red) after 15 days of Il-15 expansion. Shown also is the fluorescence minus one control (yellow). B) K562 target cell lysis by primary NK cells shown in (A). C) Control or CD56-deficient NK cells from 3 healthy donors were incubated for 1 week after CD56 CRISPR-Cas9 delivery in 25 ng/mL IL-15 then cells were isolated by FACS and cultured for an additional 8 days and the MFI of CD56 was measured by flow cytometry. D) Specific lysis of K562 target cells by isolated and expanded CD56^bright^ NK cells from the 3 healthy donors shown in (C). E) Primary NK cells were incubated and expanded for 14 days in the presence of 50 ng/ml IL-15 then cytotoxicity against K562 target cells was tested in the presence or absence of Pyk2 inhibitor PF431396. Freshly isolated, non-expanded NK cells were used as a control and similarly treated with PF431396. F) WT or CD56-KO NK92 cells were incubated for 7 days in the presence of 50 ng/ml IL-15 then cytotoxicity was tested in the presence or absence of PF431396. Shown is one representative experiment from 3 independent biological repeats. Error bars indicate technical repeats (3), SEM.

NK92 cells are phenotypically and genotypically more aligned with the CD56^bright^ NK cell subset, whereas YTS cells are more similar to the CD56^dim^ subset (49). Therefore, we reasoned that the effect of CD56 deletion may be greater on the CD56^bright^ subset, especially given its significantly higher expression of CD56. As the CD56^dim^ subset represents the majority of NK cells found within peripheral blood and has significantly decreased expression of CD56 relative to the CD56^bright^ subset, we isolated CD56^bright^ NK cells and repeated the deletion of CD56 and measurement of cytotoxic function following IL-15 expansion. While CD56^bright^ cells do not have significant lytic function when freshly isolated, brief cytokine stimulation with IL-15 is sufficient to confer substantive cytotoxic function on this subset (57). Following IL-15 expansion, both mock transfected and CD56-KO CD56^bright^ NK cells showed robust cytolytic function against K562 target cells (Fig. 6C). Therefore, despite the requirement for CD56 in NK92-mediated cytolytic function, loss of CD56 in CD56^bright^ or CD56^dim^ NK cells does not lead to a defect in primary NK cell cytotoxicity.

We hypothesized that the expansion of primary cells in IL-15 during the process of cell editing may be overcoming the requirement for Pyk2 in primary NK cells, particularly given previous reports of IL-15 signaling altering primary NK cell dependence on Pyk2 (58). Therefore, we expanded primary NK cells for 15 days in IL-15 and tested the effect of Pyk2 inhibition on cytotoxic function. While Pyk2 inhibition moderately decreased lytic function against K562 target cells following IL-15 activation, the effect was greater in cells that had not been stimulated with IL-15, suggesting that long-term culture in IL-15 can reduce dependency of primary NK cells on Pyk2 Y402 phosphorylation for lytic function (Fig. 6E).

We further hypothesized that if long-term stimulation in IL-15 could overcome Pyk2 dependency in primary cells, it may be possible to rescue the defect in CD56-KO NK92 cells with IL-15 stimulation. We incubated NK92 WT and CD56-KO cells in IL-15 for 5 days to mimic the conditions used to generate CD56-deficient NK cells. While this stimulation increased the lytic function of WT NK92 cells, activation by IL-15 failed to rescue the cytolytic deficiency in NK92 CD56-KO cells (Fig. 6F). However, NK92 cells stimulated with IL-15 also remained sensitive to Pyk2 inhibition, suggesting that the mechanism used to overcome Pyk2 dependence in primary cells in response to IL-15 was not functional in NK92 cells.

### CD56 co-localizes to the IS with pPyk2 Y402 in freshly isolated primary NK cells

Despite intact cytolytic function in CD56-deficient primary NK cells, we sought to define the localization of CD56 in primary NK cell conjugates and we performed fixed cell confocal imaging of freshly isolated ex vivo NK cells conjugated to K562 target cells. As with NK cell lines, we found that CD56 was localized to the uro-pod of migrating cells as previously described (37). However, we also frequently found redistribution of a pool of CD56 to the immune synapse (Fig. 7A). This was quantified by measurement of the mean fluorescence intensity of CD56, which demonstrated greater CD56 intensity at the immune synapse than in non-synaptic regions of the NK cell (Fig. 7B). Furthermore, CD56 and pPyk2 Y402 were spatially co-localized at the immune synapse, and intensity of pPyk2 was similarly greater at the synapse than in non-synaptic regions as previously described (Fig. 7B) (37). Therefore, as with NK cell lines, CD56 and Pyk2 colocalize in primary NK cells and are recruited to the immunological synapse.

**Fig. 7.**
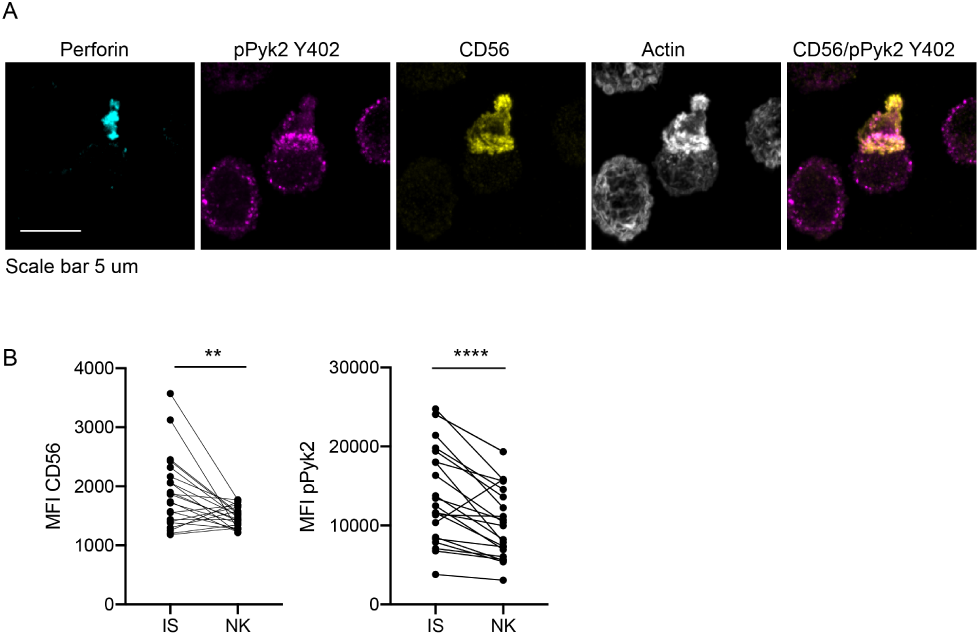
CD56 co-localizes with pPyk2 Y402 to the immune synapse in primary human NK cells. Primary NK cells were enriched from peripheral blood then incubated with K562 target cells for 45 minutes on poly-L-lysine coated coverslips in the presence of non-blocking anti-CD56 antibody. Following incubation cells were fixed, permeabilized, and immunostained for pPyk2 Y402 (magenta), perforin (cyan) and actin (phalloidin, greyscale). 3D volumetric images were acquired by spinning disk confocal microscopy. A) Representative images from one of three healthy donors. Shown is a maximum projection of 13 planes taken with 0.5 µm steps. B) Fluorescence intensity of CD56 (left) or pPyk2 Y402 (right) measured at the synaptic (IS) or non-synaptic (NK) cell cortex of primary NK cells conjugated to target cells. n=22 from one independent experiment from 3 healthy donors. **p<005, ****p<0.0001 by paired t-test.

## Discussion

While CD56 is the prototypical identifier of human NK cells in peripheral blood, its function has been poorly defined. Early studies of its role in cytotoxicity largely focused on CD56 homotypic interactions and found that cytotoxicity of IL-2 expanded primary NK cells was diminished against CD56^+^ target cells in the presence of anti-CD56 blocking antibodies (30). In contrast, Lanier et al. found no significant difference in NK cell-mediated lysis of CD56^−^ KG1a and CD56^+^ KG1a target cells (4). In both cases, CD56 functionality was tested with IL-2 expanded primary NK cells using antibody to inhibit CD56 homotypic interactions, through the manipulation of CD56 expression on target cells, or a combination of both. CD56 was also implicated in alloantigen-specific recognition by human NK cells, and the cytotoxic activity of the NS2 cell line against its LCL stimulator cell line was diminished in the presence of monoclonal antibodies specific for CD56 (59). Conversely, the expression of NCAM on some target cell lines inhibits NK cell-mediated cytotoxicity (29). Here, using CD56-deficient NK92 and YTS human NK cell lines, we demonstrate a requirement for CD56 in killing of CD56-negative targets by human NK cell lines and define a mechanism for the requirement for CD56 in human NK cell cytotoxicity.

Our previous study defined a role for CD56 in human NK cell migration, and we had previously modeled this requirement using the NK92 cell line (35). Here we extended our studies to the use of YTS cells as a second human NK cell line. Despite a significant and consistent effect of CD56 deletion on the cytotoxic function of NK92 cells, this effect was not conserved between the two human NK cell lines that we tested. The profound deficiency in lytic function in the NK92 cell line was observed even when effector cells were incubated with targets for just one hour in a ^51^Cr release assay, a timescale which likely only permits a single target cell killing to occur (41). As this incubation time led to significant killing of target cells by WT NK92 cells, these data suggest that the observed decrease in killing by the NK92 CD56-KO cells is not due to an inability to mediate serial killing against multiple targets despite the previously demonstrated requirement for CD56 in NK92 cell migration (35, 44). Furthermore, our subsequent recapitulation of cytotoxicity using antibody-coated glass to activate NK cells uncoupled the defect that we describe in exocytosis from one of cell migration.

For YTS cells, killing was only impaired at the 1-hour time point, signifying an initial delay in their ability to kill which could be rescued at later time points. The milder effect on cytotoxicity in the YTS cell line when compared to the NK92 cell line suggests fundamental differences in the requirements of the two cell lines for cytolytic function, and it remains possible that the effect on YTS cell cytotoxicity could be due to impaired cell migration affecting serial killing of targets at later time points given the requirement for CD56 in human NK cell migration (35). However, impaired IFNγ production by YTS CD56-KO cells following contact-dependent co-incubation with 721.221 target cells further underscores underlying differences in the requirement for CD56 in these cell lines and also demonstrates a functional relevance for CD56 in the YTS cell line. Previously demonstrated phenotypic and genotypic differences between the cell lines (49) supports this hypothesis and suggests that further investigation is required to fully delineate the relative role of CD56 in these contexts. Here, we have chosen to focus on the mechanism underlying the requirement for CD56 in the cytotoxic function of NK92 cells.

Our investigations into the mechanism of CD56 function in human NK cells were informed by extensive investigations into the functional role of NCAM in neural cells, where NCAM-mediated signaling plays a critical role in axonal growth, survival and proliferation (15). While there are multiple pathways by which this can occur that include homotypic and heterotypic NCAM interactions, NCAM signaling includes the formation of complexes that signal through Ras-MAPK-ERK pathways via interactions with p59fyn and FAK. Specifically, NCAM140 constitutively interacts with p59fyn, whereas FAK is recruited upon cross-linking of the NCAM extracellular domain (14). Recruitment of FAK leads to its phosphorylation and kinase function, and ultimately growth cone migration. The demonstrated requirement for Src family kinases, including Fyn, in NK cell cytotoxicity suggested that a common pathway could be functioning in NK cells. Furthermore, while FAK is not known to be expressed in human NK cells, the Src family kinase Pyk2 is expressed and is required for polarization of the MTOC during IS formation and resultant cytotoxic function (37). Here, we define a requirement for CD56 in phosphorylation of Pyk2 on Y402, the primary autophosphorylation site of Pyk2 that serves as a docking site for the SH2 domain of Src family kinases (60). This requirement was most significant in the NK92 cell line, which we found had a significantly greater amount of Pyk2 Y402 phosphorylation both at baseline and upon activation. The functional implication of this finding was the significantly greater impairment in NK92 cytotoxicity following pre-treatment of both cell lines with a Pyk2 inhibitor.

Despite showing that Pyk2 is required for NCAM-mediated function, it is still unclear precisely the mechanism by which this signaling occurs. We demonstrate that loss of CD56 leads to abolishment of lytic granule exocytosis, as surface CD10^7^a was decreased on NK92 CD56-KO cells compared WT NK92 after activation with either plate-bound antibodies or with target cells. In addition, the maximum activity of granzyme A in the supernatant, an indirect readout of exocytosis, was decreased for NK92 CD56-KO cells. Given that the exocytosis defects we observed in the NK92 cell line were demonstrated using antibody cross-linking, and thus independently of target cell activation, it is unlikely that CD56 is binding directly to unknown target cell ligands in this context. Furthermore, our use of CD56-negative target cells defines independence from homotypic NCAM interactions. NCAM can also bind both in cis and in trans to FGFR, and interactions between CD56 on NK cells and FGFR1 on T cells is sufficent to induce T cell IL-2 production (33). Despite this indication that productive interactions occur within immune cells upon CD56-FGFR1 interactions, we have repeatedly failed to detect FGFR1 on human NK cells or the target cells used in this study (data not shown), suggesting that FGFR1 is not the mechanism by which NK cell function is mediated by CD56.

Given the defect in exocytosis in CD56-deficient NK92 cells, we sought to define earlier steps in NK cell cytotoxicity and found that actin accumulation at the synapse was decreased in CD56-KO cells, although not significantly, and that MTOC polarization was impaired. Given the previously described requirement for Pyk2 in MTOC polarization during NK cell cytotoxicity (37), these data further suggest that the requirement for CD56 is in Pyk2-mediated function during cytotoxicity. This was further defined by impaired Pyk2 phosphorylation and recruitment to the IS in NK92 cells. Taken together, we propose a mechanism by which CD56 helps to localize or retain Pyk2 at sites of potential activation, where it could be available for subsequent autophosphorylation and activation. Such a model has been proposed as the mechanism by which NCAM signals through FAK (14), however there are other mechanisms that could be at play, including the previously described role for NCAM in regulating lipid raft inclusion and exclusion of signaling receptors, which was beyond the scope of this study (61).

While demonstrating a requirement for CD56 in cytotoxicity mediated by NK92 cells, we found that primary NK cells mediated target cell killing when CD56 was deleted from either total NK cells or specifically from the CD56^bright^ subset. Given the challenges inherent in manipulating gene expression in primary cells, we were reliant upon the use of cytokines, in this case IL-15, to promote NK cell survival and expansion following gene editing. It remains unclear as to whether this expansion bypasses a requirement for CD56 function in primary NK cells, although we do demonstrate that IL-15 stimulation of ex vivo NK cells significantly reduces reliance on Pyk2 Y402 phosphorylation for NK cell lytic function based upon the effect of Pyk2 inhibitor. Despite the apparent reduced dependence on Pyk2 phosphorylation in IL-15 activated primary cells and the robust phosphorylation of Pyk2 Y402 in NK92 cells, activation of the NK92 cell line with IL-15 failed to rescue the defect in cytotoxicity in NK92 CD56-KO cells. This, combined with the observation that Pyk2 inhibition does not abrogate NK92 lytic function to the same extent as CD56 deficiency, suggests that there may be other mechanisms by which CD56 is exerting function in human NK cells. Ultimately, we show that despite the requirement for CD56 function in cell lines, ex vivo human NK cells are independent of this requirement, a finding that has relevance for clinical applications of NK cell-mediated adoptive therapies.

Consistent with previous reports (4), our data demonstrate that the transmembrane-containing 140 kDa NCAM (CD56) isoform is primarily expressed on human NK cells. Western blot analyses demonstrated a molecular weight range of 130-160 kDa, which was significantly decreased following treatment of NK cell lines with PNGase F to remove polysialic acid. While these experiments, which led to the generation of a band that appeared close to 120 kDa, suggested that NCAM-120 could be expressed intracellularly (62), treatment with PI-PLC, which we demonstrated cleaved GPI-anchored proteins, did not affect the detection of CD56 on the cell surface. The single band that we detected was also consistent with the predicted weight of the NCAM140 isoform containing 858 amino acids (39). Furthermore, expression of the extracellular domain (ΔICD) or the intracellular domain (ΔECD) of CD56 failed to restore cytotoxic function to the NK92 CD56-KO, whereas expression of full-length 140 kDa NCAM was sufficient. These experiments also concur with previous evaluations of transcript level expression that also describe the predominant isoform in human PBMCs as the 140 kDa isoform (39). While other reports have suggested that NCAM120 is the primarily expressed isoform on freshly isolated NK cells based upon qPCR (63), the use of Western blotting in our study allows us to directly assess protein expression. The degree of polysialation in human NK cells is notable, however, particularly in NK92 cells and primary NK cells. The high level of polysialation in human NK cells suggests that polysialic acid itself may be playing a role in CD56 function, although further investigation is warranted.

In summary, here we describe a novel, functional role for CD56, the canonical identifier of human NK cells. We build upon previous studies describing a role for CD56 in NK cell cytotoxicity and those that define its role in cell migration and NK cell maturation to elucidate, for the first time, its role in the lysis of CD56-negative targets. In doing so we describe an unexpected role for CD56 in the function of commonly used NK cell lines, including NK92 cells, but demonstrate that primary NK cells can execute cytotoxicity independently of CD56. Finally, we link the cytotoxicity defect in CD56-KO NK92 cells to impaired signaling through the non-receptor tyrosine kinase Pyk2, and thus define novel roles for CD56 and Pyk2 in human NK cell cytotoxic function.

## Methods

### Human NK cells

For imaging experiments, primary human NK cells were isolated by venipuncture followed by NK cell enrichment using RosetteSep (Stemcell Technologies) and separation by Ficoll-Paque density gradient. Cells were resuspended in complete R10 media then co-cultured with K562 target cells and prepared for fixed cell confocal imaging as described below. Primary cells for CRISPR-Cas9 deletion experiments were obtained from leukopaks obtained from anonymous healthy platelet donors and enriched using RosetteSep and Ficoll-Paque Plus according to manufacturer’s instructions.

### Cell culture and generation of CD56-KO cell lines

NK cell lines and target cell lines were maintained at an approximate concentration of 1-5 × 10^5^ cells/ml and cultured at 37°C 5% CO2. YTS cells were a generous gift from Dr. J. Orange (Columbia University) and NK92 cells were acquired from ATCC. RPMI supplemented with 10% fetal bovine serum (FBS) plus essential nutrients was used to culture YTS cell lines and 721.221, K562 and KT86 target cells (42). NK92 cell lines were maintained in Myelocult media (Stemcell Technologies) supplemented with 200 U/ml of IL-2 (Roche) and 50 U/ml of penicillin/streptomycin (Gibco). All cell lines were routinely confirmed to be mycoplasma negative and cell line identify was routinely confirmed by flow cytometry assessing expression of known surface markers.

NCAM1 (CD56) was deleted in YTS using CRISPR gene editing as previously described for our generation of the NK92 cell line (35). In brief, a U6gRNA-Cas9-2A-GFP vector (Sigma-Aldrich) containing a NCAM1-specific gRNA sequence was incorporated into the NK cell lines through nucleofection. 10^6^ cells per condition were nucleofected with 4 µg of DNA using the Amaxa Nucleofector II (Lonza, Kit R; program R-024). Nucleofected cells were allowed to recover for 24-48 hours before sorting for GFP positivity and lack of CD56 expression.

Re-expression of CD56 in NK92 cell lines was performed by retroviral transduction. Phoenix-AMPHO cells (ATCC) were transfected with 4.5 µg of the pBabe-puro-NCAM140 mApple construct (NCBI reference sequence: NM_000615.6, transcript variant 1) using Fugene 6 Transfection Reagent (Promega). Supernatant containing viral particles was collected and concentrated using PEG-IT (System Biosciences) followed by centrifugation at 2600 rpm. NK92 CD56-KO cells were transduced with pBabe-puro-NCAM mApple viral particles in the presence of MAX Enhancer and TransDux (System Biosciences). Within 48 hours cells were sorted for CD56 and mApple and cells were expanded and maintained under antibiotic selection with regular phenotypic validation.

### Primary NK cell electroporation

Primary human NK cells were incubated after enrichment for 16 hours at 37°C in 3 ng/mL IL-15 (Miltenyi) in RPMI media supplemented with 10% Human AB serum (Sigma). Cells were washed with PBS, twice, and suspended at 2 × 10^7^ cells/ mL in EP Buffer (MaxCyte). Using OC-100 product assemblies and the MaxCyte GT (Maxcyte), cells were electroporated in the presence of Cas9 mRNA (TriLink) and CD56 guide RNA (CGCU-GAUCUCCCCCUGGCU; IDT) using the WUSTL-3 setting and transferred to a 12-well plate and allowed to rest for 10 minutes at 37°C (64). Pre-warmed media containing 3 ng/mL IL-15 was added to the cells which were then cultured in IL-15 containing media as indicated. Cells were sorted based on CD56 expression on BD FACS Aria II to >90% purity. Flow-based killing assays were performed for 1 hour using CFSE-labeled K562 as previously described (65).

### Chromium release assay

NK cell effector cells were co-cultured with target cells that had been pre-incubated with 100 µCi ^51^Cr for 1 or 4 hours in 96-well round-bottomed plates at 37°C 5% CO2. 1% IGEPAL (v/v) (Sigma-Aldrich) was used to lyse maximal release control wells and plates were centrifuged. Supernatant was transferred to a LUMA plate (Perkin Elmer) and dried overnight. Plates were read with a TopCount NXT and % specific lysis was calculated as follows: (sample – average spontaneous release) / (average total release – average spontaneous release) x 100.

### BLT esterase assay

96-well flat-bottomed plates were pre-coated overnight at 4°C with 5 µg/ml anti-NKp30 (clone P30-15, Biolegend) and -CD18 (clone IB4) or with mouse IgG1a*x* as an isotype control. Wells were blocked with phenol red-free RPMI complete medium then washed three times with PBS. 10^5^ WT or CD56-KO NK92 cells were added per well. Plates containing cells were briefly centrifuged then incubated for 90 minutes at 37°C 5% CO2. Following incubation 1% v/v IGEPAL Sigma-Aldrich) was added to maximum release wells and plates were centrifuged at 1000 rpm. Supernatant was transferred and substrate solution containing PBS, HEPES (Gibco), N-α-Cbz-L-lysine thiobenzyl ester hydrochloride (BLT; Sigma-Aldrich) and 5,5’-Dithiobis(2-nitrobenzoic acid) (DTMB; Sigma-Aldrich) was added. The plate was incubated at 37°C 5% CO2 for 30-60 minutes and luminescence was read at 415 nm using the BioTek Synergy H4 Hybrid plate reader. % maximum activity was calculated as: (sample absorbance – average background)/(average total release – average background) x 100%.

### CD107a degranulation assay and detection of phosphorylated proteins by flow cytometry

NK92 and YTS cell lines were assessed for surface CD10^7^a expression as a marker for degranulation after activation. 2 × 10^5^ WT or CD56-KO NK92 and YTS cell lines were added to 12-well flat-bottomed plates that were pre-coated overnight at 4°C with anti-NKp30 (clone P30-15, Biolegend) and -CD18 (clone IB4) (5µg/ml each) or 5% bovine serum albumin as a negative control. Alternatively, effector cells were cocultured with 10^5^ 721.221 target cells (2:1 effector to target ratio) in 5 ml polystyrene round-bottom tubes. Each condition was run in triplicate. Anti-CD10^7^a antibody (eBioscience, clone eBioH4A3) was added at the onset of incubation for co-culture experiments. Cells added to the pre-coated antibody plates were spun briefly at 200 rpm before incubation for 90 minutes at 37°C 5% CO2. Cells activated by plate-bound antibodies were mechanically dislodged and transferred to 5 ml polystyrene round-bottom tubes and incubated for an additional 25 minutes with anti-CD107a. Cells were fixed using 300-400 µl of 2% paraformaldehyde (Electron Microscopy Sciences, catalog 15712-S), then CD107a surface expression was measured using a modified LSR Fortessa (BD Biosciences) with the CD107a+ gate defined by unstained negative control cells. Data were analyzed using FlowJo X (TreeStar Inc.) software. For experiments performed with target cell activation the average background of media alone was subtracted from samples. For detection of phospho-Pyk2, cells were activated as above for and then fixed and permeabilized followed by detection with pPyk2 Y402 (Abcam, ab4800) and secondary detection with goat anti-rabbit IgG Alexa Fluor 488 (Invitrogen). Mean fluorescence intensity was adjusted to relative fluorescent intensity by normalizing to the WT condition.

### Flow cytometry

FACS analysis of WT or CD56-KO NK92 and YTS cell lines was performed using modified NK cell panels designed to assess the expression of adhesion, inhibitory, and developmental ligands and intracellular molecules (Table 1) (47). For panels evaluating response to activation 10 ng/ml phorbol 12-myristate 13-acetate (PMA, Sigma-Aldrich) and 1 µg/ml of ionomycin were used to stimulate cells for 4 hours at 37°C 5% CO2. 10 µg/ml of Brefeldin A (Sigma-Aldrich) was added at the onset of incubation to stimulated and unstimulated controls to prevent protein transport of newly synthesized proteins in response to cellular activation. After incubation, activated cells were stained for surface ligands for 25 minutes followed by permeabilization and fixation using BD Cytofix/Cytoperm (BD Biosciences). Antibody staining was performed in the dark and at room temperature; cells were then stained for intracellular markers for 1 hour before being washed and resuspended in PBS 1% paraformaldehyde (Sigma-Aldrich). Cells in panels without PMA/ionomycin stimulation were also resuspended in PBS 1% paraformaldehyde after being incubated with antibodies for 25 minutes. Detection of CD56 on cell lines was performed using clone HCD56 (1:100, Biolegend). For evaluation of cell line CD56 expression, detection of polysialated NCAM (PSA-NCAM) was by clone 12F8 (BD Biosciences) followed by anti-rat goat IgG FITC (Invitrogen). 10^4^ events per sample were acquired using a modified LSR Fortessa (BD Biosciences). Fluorescence minus one controls (FMO) were used to define positive and negative populations. Prism 6.0 (GraphPad Software) was used to graph the percent positive and the mean fluorescence intensity calculated using FlowJo X (TreeStar Inc.) software.

**Table 1:** Antibodies used for flow cytometric evaluation of cell lines.

| Antibody | Clone | Fluorochrome | Antibody Source | Host | Antibody dilution |
| --- | --- | --- | --- | --- | --- |
| NKp46 | 9E2 | Pacific Blue | Biologend | Mouse | 4/85 |
| CD56 | HCD56 | Brilliant Violet 605 | Biologend | Mouse | 1/85 |
| CD16 | 3G8 | Brilliant Violet 650 | Biologend | Mouse | 1/340 |
| DNAM-1 | 11A8 | FITC | Biologend | Mouse | 1/17 |
| NKp44 | Z231 | PE | Beckman Coulter | Mouse | 1/17 |
| NKG2D | 1D11 | PE-Cy7 | Biologend | Mouse | 1/68 |
| CD69 | FN50 | PE-CF594 | BD Biosciences | Mouse | 1/68 |
| NKp30 | P30-15 | APC | Biologend | Mouse | 1/85 |
| CD16 | B73.1 | APC Cy7 | BD Biosciences | Mouse | 1/17 |
| CD25 | B1.49.9 | APC-Alexa Fluor 700 | Beckman Coulter | Mouse | 1/85 |
| CD2 | 39C1.5 | Pacific Blue | Beckman Coulter | Rat | 1/34 |
| CD244 | 2-69 | FITC | BD Biosciences | Mouse | 1/17 |
| CD11c | BU15 | PerCP Cy5.5 | Beckman Coulter | Mouse | 1/34 |
| CD28 | L293 | PE | BD Biosciences | Mouse | 1/17 |
| CD11a | HI111 | PE-Cy7 | BD Biosciences | Mouse | 1/85 |
| CD18 | 6.7 | APC | BD Biosciences | Mouse | 1/17 |
| CD158e | DX9 | Brilliant Violet 421 | Biologend | Mouse | 3/64 |
| CD158b | DX27 | FITC | Biologend | Mouse | 1/32 |
| CD94 | HP-3D9 | PerCP Cy5.5 | BD Biosciences | Mouse | 1/80 |
| NKG2C | 134591 | PE | R&D Systems | Mouse | 1/32 |
| CD158a/h/g | HP-MA4 | PE-Cy7 | Affymetrix eBioscience | Mouse | 1/32 |
| KIR2DS4 | 179315 | APC | R&D Systems | Mouse | 1/40 |
| NKG2A | 131411 | Alexa Fluor 700 | R&D Systems | Mouse | 1/80 |
| CD57 | NC1 | Pacific Blue | Beckman Coulter | Mouse | 1/160 |
| CD62L | DREG-56 | Brilliant Violet 650 | BioLegend | Mouse | 1/160 |
| CD127 | A019D5 | Brilliant Violet 785 | BioLegend | Mouse | 1/64 |
| CD122 | TU27 | PE | BioLegend | Mouse | 1/80 |
| CD117 | 104D2D1 | PE-Cy7 | Beckman Coulter | Mouse | 1/16 |
| CD27 | O323 | PE-Cy5 | Affymetrix eBioscience | Mouse | 1/64 |
| CD94 | DX22 | APC | Biologend | Mouse | 1/80 |
| CD11b | Bear1 | APC-Alexa Fluor 750 | Beckman Coulter | Mouse | 1/32 |
| Perforin | B-D48 | Brilliant Violet 421 | Biologend | Mouse | 1/80 |
| TNFA | MAb11 | Brilliant Violet 650 | Biologend | Mouse | 1/32 |
| Perforin | dG9 | FITC | BD Pharmingen | Mouse | 1/20 |
| CD107a | eBioH4A3 | PE-Cy5 | Affymetrix eBioscience | Mouse | 1/400 |
| Granzyme B | GB11 | PE Texas-Red | ThermoFisher Scientific | Mouse | 1/64 |
| IFNg | 4S.B3 | Alexa Fluor 700 | Biologend | Mouse | 1/32 |
| Eomes | WD-1928 | eFluor 660 | ThermoFisher Scientific | Mouse | 1/100 |
| T-Bet | 4B10 | PE | Biologend | Mouse | 1/9 |

### IFNγ detection by ELISA

WT YTS and CD56-KO cells were incubated with 721.221 target cells at a 2:1 ratio at 37°C 5% CO2 in round-bottomed 96-well plates for 22 hours in triplicate. Plates were centrifuged and the supernatant collected. Human IFNγ was detected by ELISA (Abcam) following the manufacturer’s procedure. Absorbance was read at 415 nm on a BioTek Synergy H4 Hybrid plate reader. IFNγ concentrations were calculated using a standard curve following subtraction of background (media only) from all conditions. Results were graphed and statistical analyses were performed using Prism 6.0 (GraphPad software).

### PI-PLC cleavage of GPI-anchored proteins and detection by flow cytometry

NK92 and YTS NK cells, or Raji and Jurkat cells as a positive control were incubated with 1 U/ml of phosphatidylinositol-specific phospholipase C (PI-PLC; Invitrogen) at 4°C for 30 minutes in cold PBS. Cells were then washed twice with cold PBS and transferred to polystyrene tubes. Cells were immunostained for 25 minutes at room temperature with anti-CD56 (BV421; clone HCD56, Biolegend) or anti-CD55 (PE; clone JS11, Biolegend). Cells were washed once with PBS and then resuspended in 2% paraformaldehyde and analyzed by flow cytometry. Prism (GraphPad Software) was used to graph the percent positive and the mean fluorescence intensity calculated using FlowJo X (TreeStar Inc.) software.

### Lysate preparation and Western blots

Cell lysates from 5-10 × 10^6^ cells were generated using CHAPS Cell Extract Buffer (Cell Signaling Technology) supplemented with 1X Halt protease inhibitor cocktail (Thermo Fisher Scientific). Samples were incubated at 95°C with NuPAGE sample reducing agent (Thermo Fisher Scientific) and 4X NuPAGE LDS sample buffer (diluted to 1X) for 10 minutes. 2-4 × 10^5^ cell equivalents per well were loaded into a NuPAGE 4-12% Bis-Tris density gradient gel (Thermo Fisher Scientific) and ran at a constant 150V for 80 minutes. Separated proteins were transferred onto nitrocellulose membranes using a Mini Gel Tank/Mini Blot Module (Life Technologies) at a constant 0.2A for 105 minutes. The nitrocellulose membranes were then blocked with 5% nonfat milk in PBS 0.05% Tween-20 for 90-120 minutes at 4°C. Nitrocellulose membranes were incubated overnight at 4°C with primary antibodies in 5%(w/v) BSA in PBS 0.05% Tween 20 at the following dilutions: 1:1000 anti-CD56 (mouse monoclonal, clone 123C3; Cell Signaling Technology) and 1:4000 anti-actin (rabbit polyclonal; Sigma-Aldrich) as a loading control. Detection of polysialated NCAM (PSA-NCAM) was by clone 2-2B (Millipore). Membranes were washed with 0.5M NaCl in PBS 0.05% Tween 20. Primary antibodies were probed with either IRDye 680RD Goat anti-mouse IgG or IRDye 800CW goat anti-rabbit IgG (Li-COR Biosciences) secondary antibodies for 1 hour at room temperature. Nitrocellulose membranes were imaged using the Odyssey CLx imaging system (Li-COR Biosciences). Image Studio Lite software was used for densitometry and analysis of Western blot images.

### Confocal microscopy

For fixed cell imaging, WT and CD56-KO NK92 cells were co-cultured with K562 target cells at a 2:1 effector to target ratio in complete R10 medium. Cells were incubated for 20 minutes at 37°C 5% CO2 then were transferred to poly-L-lysine coated 1.5 coverslips for an additional 25 minutes. Following incubation, cells were fixed and permeabilized with CytoFix/CytoPerm (BD Biosciences) at room temperature for 15 minutes and were washed twice with 50-100 µl of PBS. Conjugate immunostaining was performed with biotinylated monoclonal mouse anti-tubulin (Invitrogen) and Brilliant Violet 421-conjugated streptavidin (Invitrogen); Alexa Fluor 488-conjugated mouse anti-perforin (clone δG9); phalloidin Alexa Fluor 568; pPyk2 Y402 (Abcam, ab4800) and goat anti-rabbit IgG Alexa Fluor 488 (Invitrogen). Where indicated, cells were incubated with anti-CD56 Alexa Fluor 647 (HCD56, Biolegend) during target cell incubation. Coverslips were mounted to slides using ProLong Gold antifade reagent (ThermoFisher Scientific). For detection of CD56 in conjugates, NK cells were pre-incubated with HCD56 Alexa Fluor 647 (Biolegend) prior to conjugate formation.

### Image analyses

Fiji (66), Volocity (Perkin Elmer) or Imaris (Bitplane) were used to process and analyze confocal image sequences. MTOC polarization to the synapse was determined using the line measurement tool after denoting the highest point of fluorescence intensity of α-tubulin as the MTOC. For actin accumulation at the synapse, the stamp tool was used to measure the fluorescence intensity of actin at the synapse and distal region of both the effector and target cells (67). The background and initial synapse intensity (AU) of actin was calculated as the area (µm^2^) x the mean of fluorescence intensity. Total intensity of actin at the synapse was calculated as follows: Total synapse intensity = synapse intensity – (effector background intensity + target background intensity). Lytic granule convergence was calculated by measuring the distance from individual granules to the MTOC as defined as the brightest point of α-tubulin intensity (54). For the accumulation of CD56 at the synapse of primary NK cells, masks of synaptic vs. non-synaptic actin were generated following auto thresholding of intensity in the actin channel. Intensity of pPyk2 Y402 or CD56 staining was measured in Fiji following auto thresholding, with the “limit to threshold” box checked. Intensity of respective channels of interest at the synapse or the non-synaptic cortical region were plotted for individual cells.

### Statistical analyses

Prism 6.0 (GraphPad software) was used for statistical analyses. Unless otherwise stated, all analyses used either a Student t-test with Welch’s correction or a one-way ANOVA corrected for multiple comparisons. Statistical significance was denoted when differences in measurements produced a p-value of < 0.05.

## ACKNOWLEDGEMENTS

This work was supported in part by NIH-NIAID R01AI137073 to EMM.

---

**Supplementary Figure 1.**
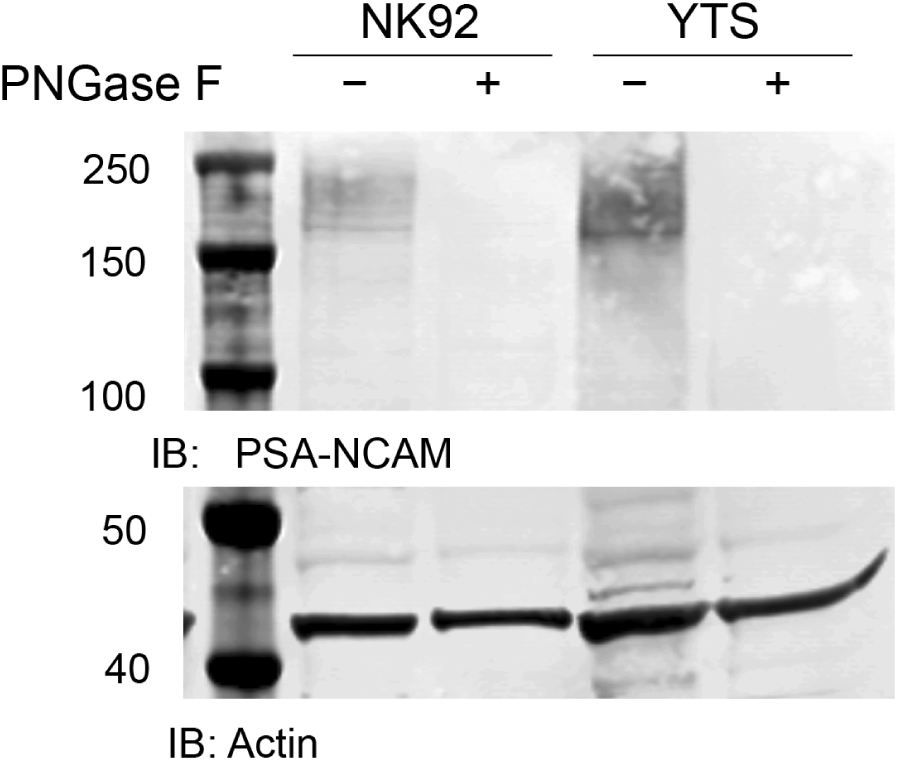
Expression of PSA-NCAM on human NK cell lines. A) NK92 or YTS cell line lysates were pre-treated with PNGase F to remove PSA-NCAM through cleavage of N-linked glycans. Following enzyme treatment, lysates from treated and untreated conditions were separated by SDS-PAGE and Western blotting with anti-PSA-NCAM antibody or actin as a loading control was performed.

**Supplementary Figure 2.**
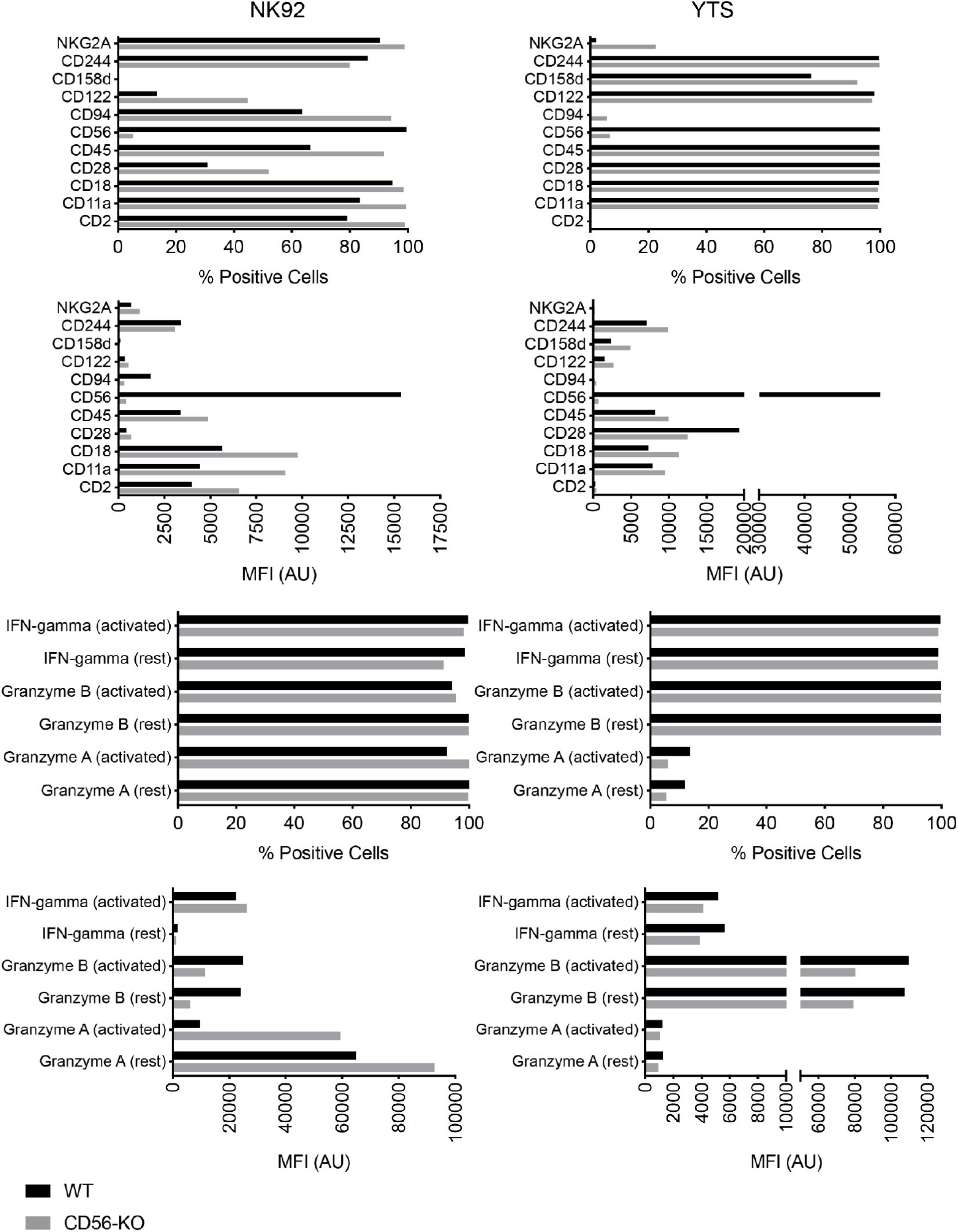
Phenotyping of NK cell lines by flow cytometry. NK92 and YTS WT and CD56-KO cells were analyzed for expression of cell surface receptors and intracellular effector molecules using five panels as described in Methods. Effector functions were evaluated in the presence or absence of activation by PMAand ionomycin. Mean fluorescence intensity was measured and % positive cells based on fluorescence minus one gating was also calculated. Shown is one representative of 3 independent experiments.

**Supplementary Figure 3.**
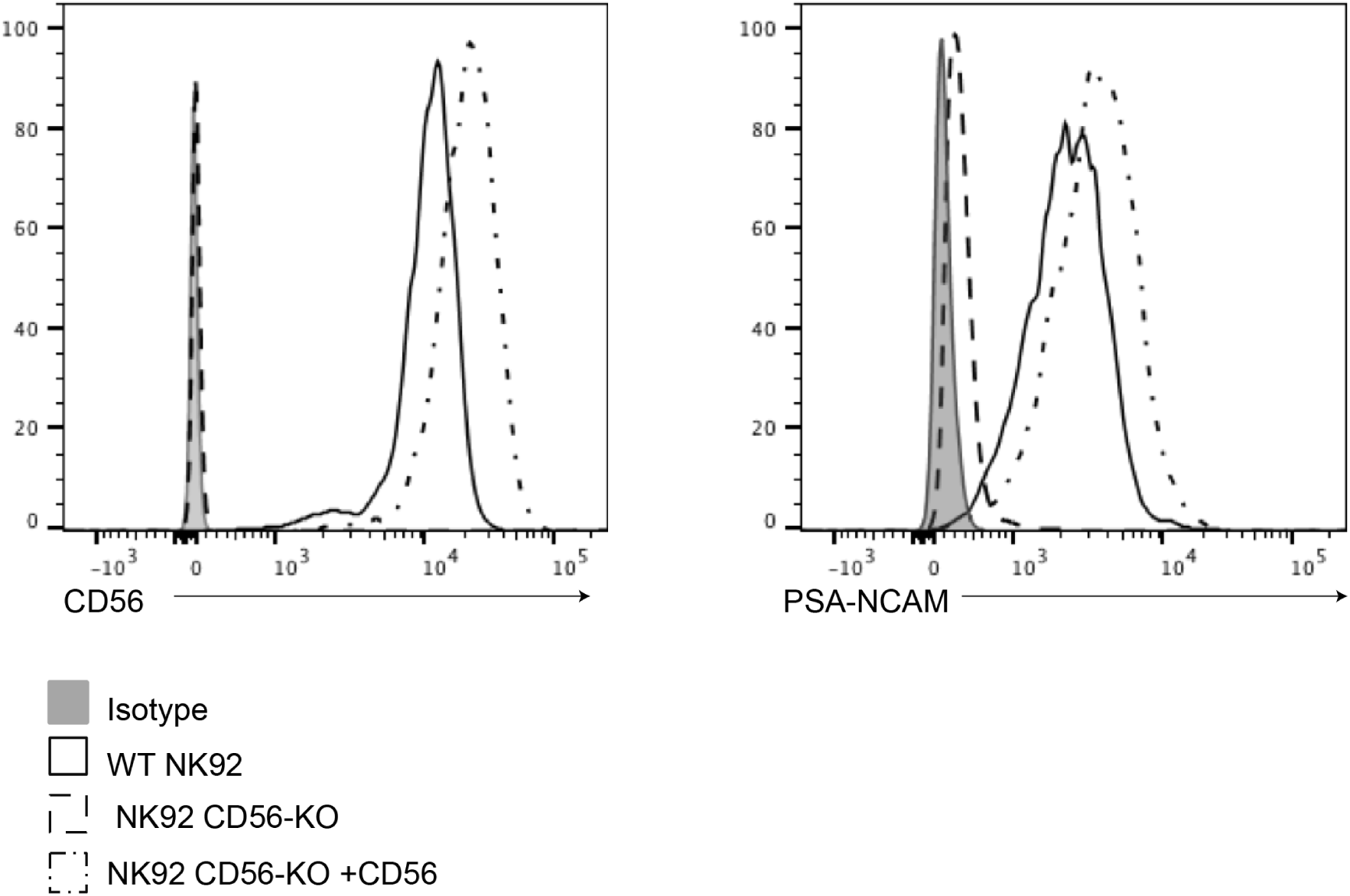
Re-expression of CD56 and PSA-NCAM in reconstituted NK92 cell lines. CD56 (left) or PSA-NCAM (right) were detected by flow cytometry on WT (solid line), CD56-KO (dashed line) or CD56-KO cells reconstituted with NCAM-140 (dot-dashed line). Isotype antibody was used as a negative control (solid filled histogram). Shown is one representative of two independent repeats.

**Supplementary Figure 4.**
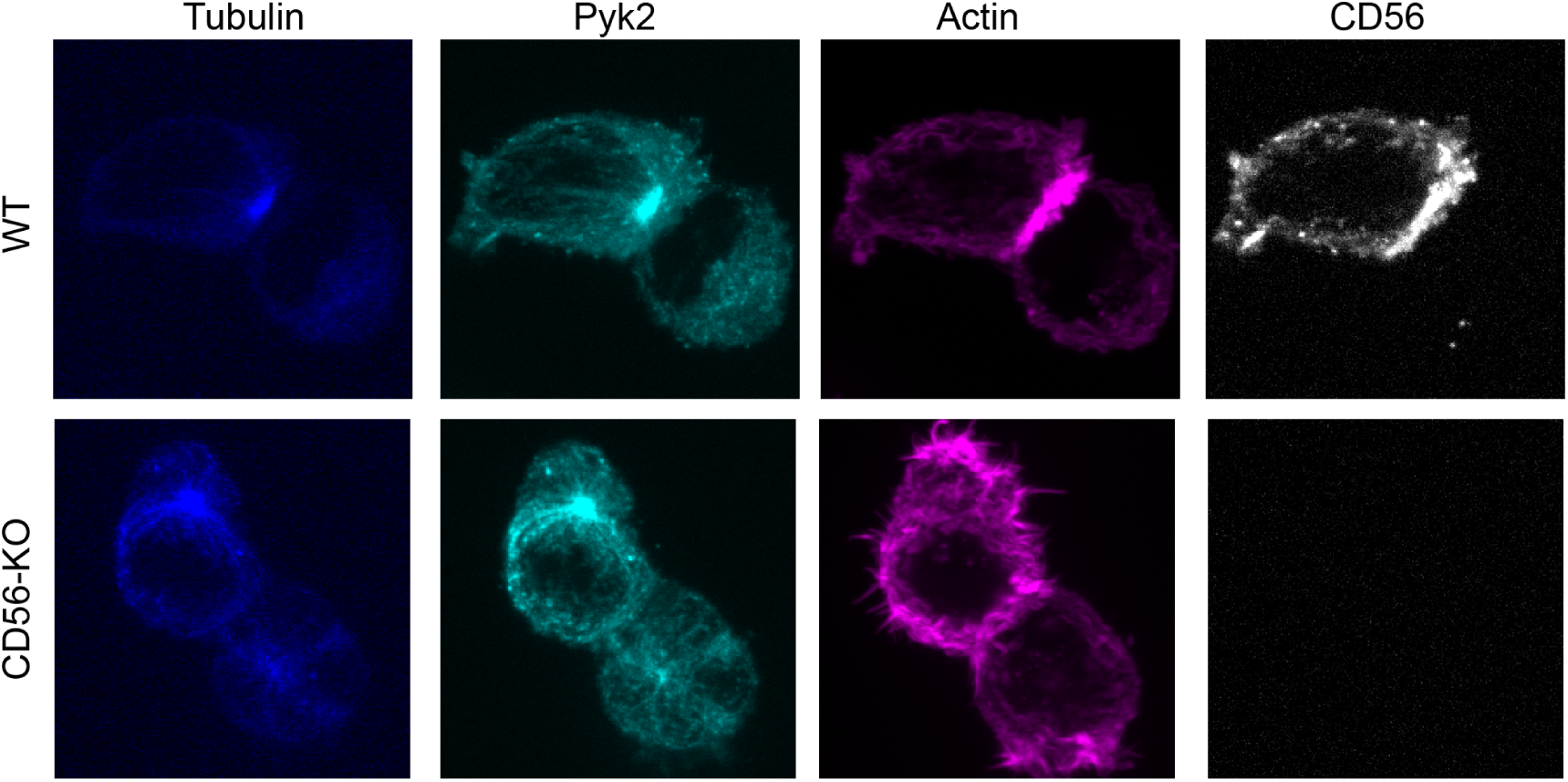
Detection of total Pyk2 in WT and CD56-KO cell lines. WT (top) or CD56-KO (bottom) NK92 cells were conjugated to K562 targets then fixed, permeabliized and immunostained for tubulin, Pyk2, actin (phalloidin) and CD56. Images were acquired on a spinning disk confocal microscope. Shown is one representative of 60 cells from two independent technical replicates.

